# A fungal pathogen manipulates phytocytokine signaling for plant infection

**DOI:** 10.1101/2023.07.04.547642

**Authors:** Chenlei Hua, Lisha Zhang, Annick Stintzi, Andreas Schaller, Hui-Shan Guo, Thorsten Nürnberger

## Abstract

Phytocytokines are hormone-like plant peptides that modulate immune homeostasis and development. Phytosulfokine (PSK) mediates plant growth and attenuates activation of plant pattern-triggered immunity (PTI). We show that the small cysteine-containing effector VdSCP8 from Verticillium dahliae (Vd) is a virulence-promoting protein that suppresses PTI in Arabidopsis thaliana and Nicotiana benthamiana. Apoplastic SCP8 suppresses immune activation through leucine-rich repeat ectodomain pattern recognition receptors. SCP8 virulence and immunosuppressive activities require PHYTOSULFOKINE RECEPTOR 1 (PSKR1), which binds PSK and forms a complex with co-receptor BAK1 for PTI suppression. We find that PSK, like SCP8, suppresses PTI, SCP8 stimulates PSKR1-BAK1 complex formation, and that Vd requires PSK signaling for host infection. SCP8 interacts with an apoplastic subtilase, and co-expression of SCP8 and subtilase inhibitors reduces PTI suppression. Our findings suggest that a multi-host plant pathogen manipulates PTI by enhancing immunosuppressive PSK signaling, likely through plant subtilase activity.

---

In nature, the interaction between plants and microbes plays a crucial role in shaping the dynamics and stability of ecosystems. While many microbial species form mutually beneficial symbioses with plants, phytopathogenic microbes use plants as substrates to complete their life cycle, causing significant damage to agricultural crops and natural plant habitats. To successfully colonize plant hosts, these microbes employ large sets of effectors to divert plant nutrients for their own reproduction and to evade or suppress host immune responses^1^.

To cope with microbial attacks, plants have evolved pattern-recognition receptors (PRR), which are located on the cell surface and activate immunity. PRRs sense conserved microbial patterns, resulting in pattern-triggered immunity (PTI)^2^. Upon ligand binding, PRRs recruit membrane-localized co-receptors, e.g. somatic embryogenesis receptor kinases (SERKs), and transduce extracellular signals to downstream signaling components including receptor-like cytoplasmic kinases^3^, mitogen-activated protein kinases^4^ and transcription factors leading to the expression of defense genes^5^. PTI responses are activated within minutes of PRR activation and prime plants for robust immune activation.

While plant pathogenic microbes utilize effectors to target host immune signaling pathways from PRR activation to defense gene activation, plants harbor intracellular immune receptors to detect potentially harmful effector activities upon delivery into host plants. Subsequently, effector-triggered immunity (ETI) in concert with early PTI activation is assumed to provide robust immunity against microbial infections.

In uninfected plants, PTI activation is under the negative control of at least two phytocytokines^6-12^. Rapid alkalinization factor (RALF), recognized by the malectin domain-containing receptor kinase FERONIA^13^, and phytosulfokine (PSK), a tyrosine-sulfated pentapeptide recognized by the leucine-rich repeat (LRR) domain-containing receptor kinase PSKR1^14^, have been implicated in PTI suppression. Several fungal plant pathogens produce and secrete RALF-like peptides to suppress PTI^15^, suggesting that manipulation of negative regulatory mechanisms of host defenses aids in host infection.

*Verticillium dahliae* (*Vd*) is a soil-borne fungus with a broad host range, including the model plants *Nicotiana benthamiana*^16^ and *Arabidopsis thaliana* (hereafter Arabidopsis). The *Vd*-derived patterns chitin, nlp20, and pg9 activate PTI in *Arabidopsis*^17-20^ via the PRR-complexes LYK5/CERK1, RLP23/SOBIR1, and RLP42/SOBIR1, respectively. Likewise, several *Vd* effectors suppress chitin-induced plant immunity^21-23^. RNAseq analyses in *Vd*-infected *N. benthamiana* plants revealed that genes encoding PSK and its receptors PSKR1 and PSKR2 are highly up-regulated upon fungal infection (Fig. 1A and Table S1). We therefore hypothesized that still unknown *Vd* effectors may engage PSK signaling to manipulate PTI in order to suppress host immunity and to establish infection. In support of this hypothesis we confirmed that *Vd* relies on PSK signaling for host plant infection. Inoculation with *Vd* spores of Arabidopsis genotypes Col-0, the PSKR1 mutant *pskr1-3*^24^, and the *serk3 serk4* double mutant *bak1-5/bkk1*^25^ revealed that the *pskr1-3* is highly resistant to *Vd* infection as compared to the wild type Col-0 and *bak1-5/bkk1* (Fig. 1B).

**Fig. 1.**
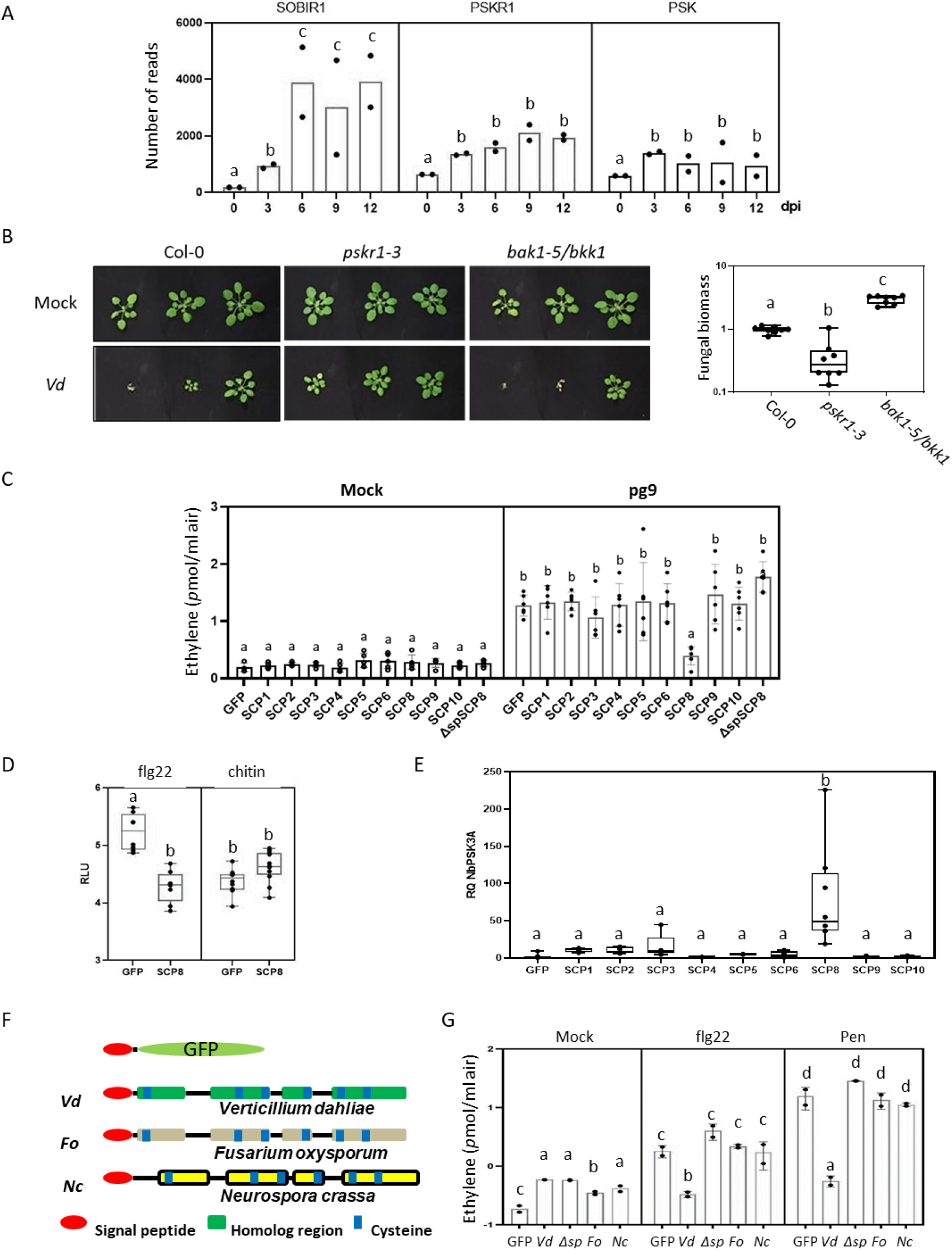
LRR-RK/RP-activated plant signalings influenced by *Verticillium dahliae* (*Vd*) and secreted cysteine containing proteins. A, Expression of *Nicotiana benthamiana PSK6, PSKR1*, and *SOBIR1* genes upon *Vd* infection at different time points. B, Susceptibility of Arabidopsis Col-0, *pskr1-3* and *bak1-5/bkk1* to *Vd* infection. Relative fungal biomass in plants was quantified by RT-qPCR analysis. C, Ethylene production of *N. benthamiana* transgenic RLP42 line transiently expressing *VdSCP* genes upon pg9 treatment. Ethylene was measured 2 hours after pg9 treatment. D, ROS burst of *N. benthamiana* transiently expressing GFP or SCP8 genes upon flg22 and chitin treatments. E, *PSK* gene (NbPSK3A) expression in *N. benthamiana* transiently expressing *VdSCP* genes. Gene expression was analysed 2 days after agro-infiltration of *VdSCP* genes by RT-qPCR. F, Structures of SCP8 from *Verticillium dahliae* (*Vd*), *Fusarium oxysporium* (*Fo*), and *Neurospora crassa* (*Nc*) with PR1 signal peptide. G, Ethylene production of *N. benthamiana* transiently expressing GFP (GFP), VdSCP8 (*Vd*), VdSCP8 without signal peptide (*Δsp*), FoSCP8 (*Fo*), and NcSCP8 (*Nc*) upon flg22 and Pen treatments. Ethylene was measured 2 hours after elicitor treatments. The different letters depict statistically significant differences (P<0.05, student’s t-test).

In previous work, we have identified a number of secreted small cysteine-containing protein effectors (VdSCP, hereafter SCP) from *Vd* that either suppressed or activated plant immunity-associated responses^26,27^. While some SCPs localized to the nucleus to target Arabidopsis transcription factors, effectors SCP8, SCP9 and SCP10 were found to accumulate extracellularly at the plant-pathogen interface^28^. To study possible immune-modulatory activities of some SCP effectors and their sites of action, we produced N-terminal fusions of the Arabidopsis *PR1* signal peptide with SCP-coding sequences for *35S*-promoter-driven transient overexpression in RLP42-transgenic *N. benthamiana* plants.

RLP42 carries a leucine-rich repeat ectodomain to recognize the fungal polygalacturonase pattern pg9, but lacks an intracellular kinase domain. We found that among all tested SCPs, only expression of SCP8, a 194-amino acid protein with 6 cysteine residues^27^, resulted in a strong reduction of ethylene production in leaves treated with pg9 (Fig. 1C). However, intracellular transient over-expression of SCP8 did not reduce pg9-induced ethylene production (Fig. 1C), suggesting that apoplastic localization of the effector is a requirement for its immunosuppressive activity. Further analysis showed that SCP8 also suppressed the flg22 (recognized by the LRR receptor kinase FLS2)-induced ROS burst. Notably, fungal chitin-induced ROS burst, mediated by its binding to the lysin-motif ectodomains of receptor kinases LYK5/CERK1 was not affected by SCP8 over-expression (Fig. 1D). These results suggest that SCP8 specifically suppresses plant immunity activated by LRR-type PRRs. We further observed that SCP8-expressing *N. benthamiana* leaves showed significant up-regulation of *PSK* gene (*NbPSK3A*) expression when compared to GFP-expressing control plants (Fig. 1E). This is in agreement with our hypothesis that SCP8 activates PSK signaling to suppress plant immunity mediated preferably by LRR-type PRRs.

*SCP8*-related genes are widespread in Ascomycota, including pathogenic fungi such as *Fusarium oxysporum* and *Magnaporthe oryzae*, and non-pathogenic fungi such as *Neurospora crassa*. Alignments of deduced amino acid sequences from pathogenic and non-pathogenic fungi reveal six conserved cysteines as well as four domain blocks (Fig. 1F). In addition, SCP8 appears to be distantly related to *Vd* elicitor PevD1^29^ and to *Alternaria alternata* major allergen Alt a 1^30^. To test whether PTI-suppressive functions are conserved among SCP8-like proteins in other fungi, *F. oxysporum FoSCP8* and *N. crassa NcSCP8*-encoding sequences were transiently expressed in *N. benthamiana* leaves for ethylene measurement upon PAMP treatment. SCP8 lacking a plant signal peptide for protein secretion was also expressed to determine functional localization of SCP8. Two PAMPs, flg22^31^ and Pen^32^, were used to treat *N. benthamiana* leaves 3 days after agro-infiltration, followed by ethylene measurements 2 hrs after the addition of PAMPs. While GFP-expressing control leaves showed significant ethylene production within 2 hours after treatment with 2 µM flg22 or 200 µg/ml Pen, VdSCP8-expressing leaves did not respond to either flg22 or Pen. Leaves expressing *FoSCP8* and *NcSCP8* (Fig 1G) showed no reduction in ethylene production. SCP8 lacking the signal peptide also failed to reduce ethylene production, suggesting that SCP8 functions as an extracellular fungal effector that evolved for a *Vd*-specific function in PTI suppression from the apoplastic space.

To assess the contribution of *SCP8* to fungal virulence, *SCP8* knock-out mutants were generated by homologous recombination^33^. Two independent gene-knockout mutants, *Δscp8-1* and *Δscp8-2* without obvious defects in fungal development (Fig. S1) were selected for host infection assays. Further, we transformed a *SCP8-GFP* gene fusion driven by either the native or the constitutive OliC promoter^34^ into the *Δscp8-1* mutant to generate the genetically complemented strains C-native and C-olic, respectively. The *Vd* wild type strain (V592), the SCP8 mutants *Δscp8-1* and *Δscp8-2*, and the complementation strains C-native and C-olic were inoculated into *Vd* host plants cotton (*Gossypium hirsutum*), Arabidopsis and *N. benthamiana*. V592 infected cotton showed severe wilt disease symptom at 30 dpi, while *Δscp8-1-* or *Δscp8-2-*inoculated cotton plants showed significantly reduced disease progression at the same time point (Fig. 2A). Similar results were also obtained for Arabidopsis and *N. benthamiana* (Fig. 2B&C). Complementation with both, native and constitutive promoter-driven *SCP8* gene constructs, recovered the virulence of *Δscp8-1* on cotton host (Fig. 2A). Likewise, two Arabidopsis transgenic lines stably over-expressing Arabidopsis *PR1 signal peptide-SCP8-GFP* constructs (T-6 and T-12) at similar levels were found to be more susceptible to *Vd* infection than wild type Col-0 (Fig. 2D). Altogether, these results demonstrate that SCP8 is required for full virulence of *Vd*.

**Fig. 2.**
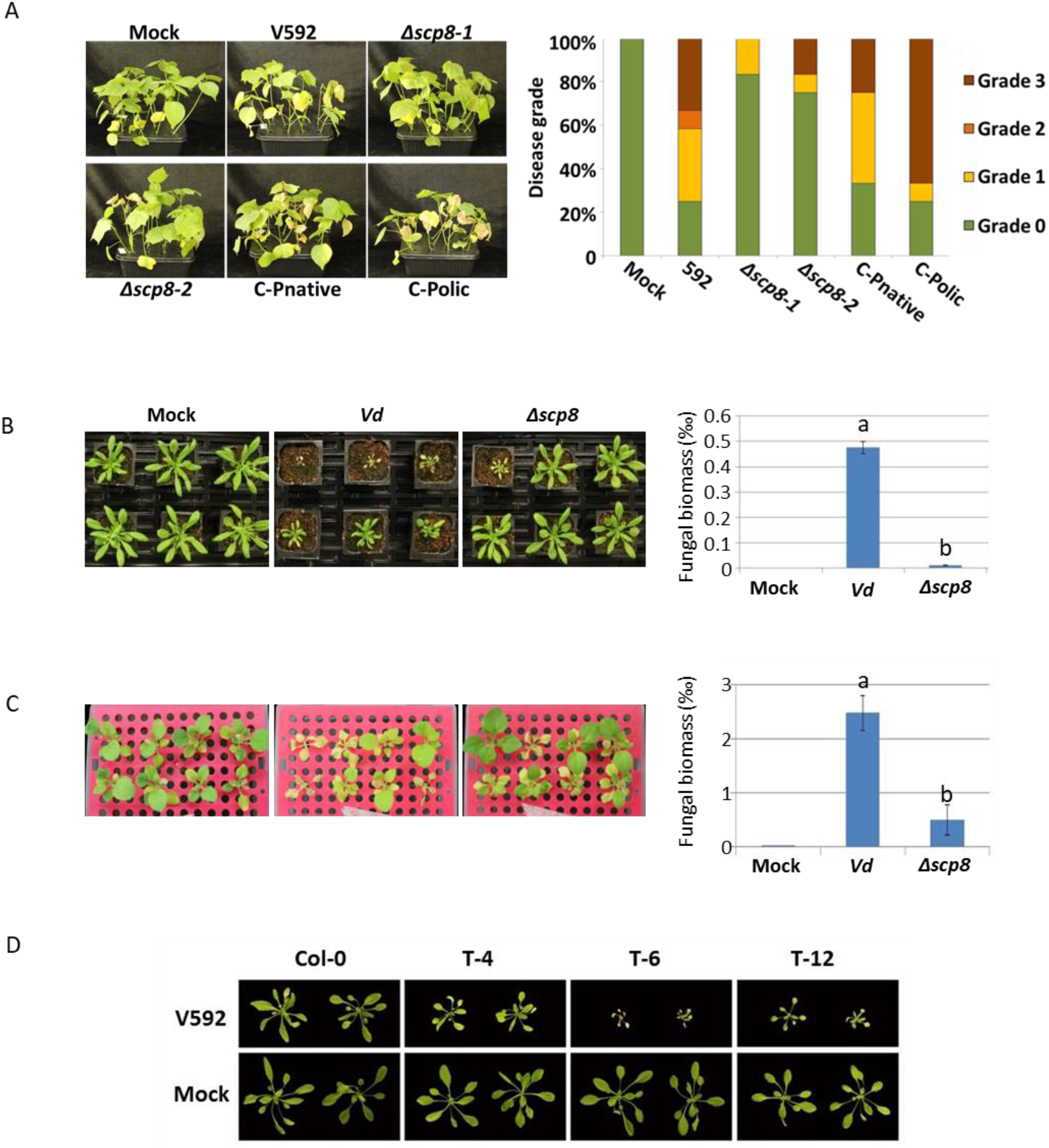
*Vd* infection assays on different plants. A, Cotton phenotypes at 20 days after inoculation of *Vd* wild-type strain V592, SCP8 gene knockout strains ⊿scp8-1 and ⊿scp8-2, gene complementary transformants C-Pnative and C-Polic. Cotton susceptibility was quantified by the ratio of plants in each disease grade following the method of Zhang *et al*. (New Phyto, 2017). B, Arabidopsis phenotypes at 20 days after inoculation of V592 and *⊿scp8-1*. Arabidopsis susceptibility was quantified by relative fungal biomass in the infected plants following the method of Fradin *et al*. (Plant Physiol. 2009). C, *N. benthamiana* phenotypes at 20 days after inoculation of V592 and *⊿scp8-1*. Arabidopsis susceptibility was quantified by relative fungal biomass in the infected plants following the method of Fradin *et al*. (Plant Physiol. 2009). D, Col-0 and transgenic Arabidopsis T-4, T-6 and T-12 expressing SCP8 infected by V592 strain. The photos of infected plants were taken at 20 days after *Vd* inoculation. The different letters depict statistically significant differences (P<0.05, student’s t-test).

Next, we treated leaves of Col-0 and transgenic lines T-6 and T-12 with flg22, nlp20, and chitin, followed by measurements of immune responses including ethylene production, ROS burst and MAPK activation. As shown in Fig 3A, the ethylene production triggered by nlp20 and pg9, but not flg22 was lower in T-6 and T-12 transgenic lines than that in wild-type Col-0 at 3 hpi. Similarly, the ROS burst of transgenic plants was significantly reduced compared to that in wild-type Col-0 in response to nlp20 or pg9 but not to chitin (Fig. 3B). Also, MAPK activation by nlp20 and flg22 was much reduced in SCP8 transgenic lines compared to wild type (Fig. 3C).

**Fig.3.**
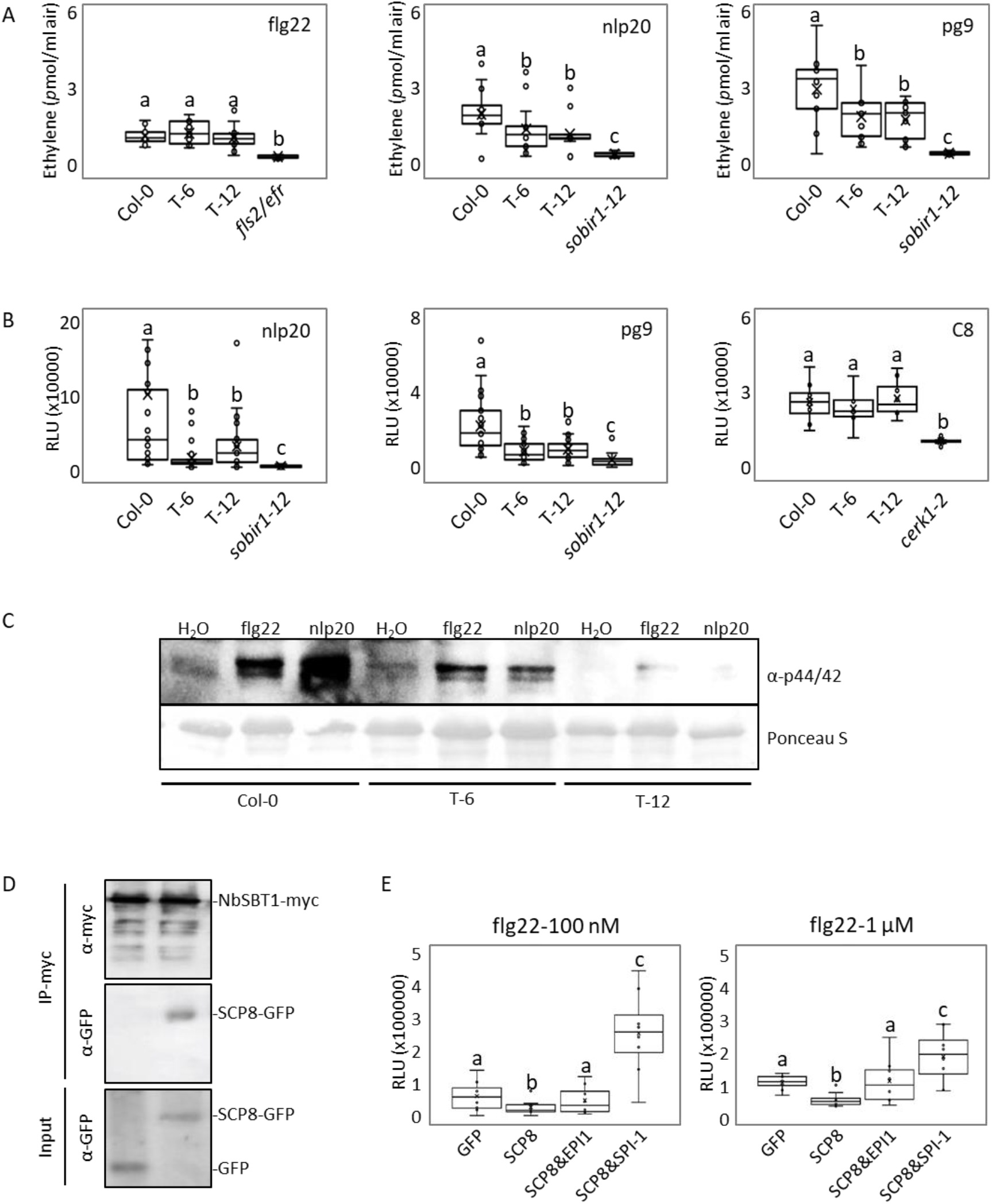
SCP8 specifically suppresses PTI activated by LRR-domain containing receptors and requires plant subtilases. A, Ethylene production of Col-0, T-6, and T-12 upon flg22, nlp20 and pg9 treatments. B, ROS burst of Col-0, T-6 and T-12 upon nlp20, pg9 and chitin (C8) treatments. C, MAPK activation of Col-0, T-6 and T-12 upon flg22 and nlp20 treatments. D, Co-immunoprecipitation and western blot in *N. benthamiana* transiently co-expressing GFP/SCP8-GFP and NbSBT1-myc. Total proteins were isolated from *N. benthamiana* leaves 2 days after agro-infiltration and immunoprecipitaed with anti-GFP agarose beads. Total proteins and immunoprecipitated proteins were detected by both anti-GFP and anti-myc antibodies. E, ROS burst of *N. benthamiana* leaves transiently expressing GFP, SCP8, SCP8&EPI1, and SCP8&SPI-1, respectively, upon different doses of flg22 treatment. The different letters depict statistically significant differences (P<0.05, student’s t-test).

To investigate the mode of SCP8 action in suppressing plant defenses, a mass spectrometry (MS)-based analysis of SCP8-interacting proteins was performed in *N. benthamiana*. Proteins were isolated and immunoprecipitated by anti-GFP agarose beads from *N. benthamiana* leaves expressing SCP8-GFP and free GFP, respectively, followed by MS analysis. Most of the peptides detected specifically in the SCP8-GFP sample match the subtilase (SBT) Niben101Scf00726g02006.1, closely related to tomato SBT3^35^ and Arabidopsis SBT1 and SBT3 families^36,37^ (Table S2). Plant SBTs are known to be involved in propeptide processing for the formation of phytocytokines, such as systemin and PSK^38,39^. Complex formation of SCP8 and *N. benthamiana* SBT was confirmed by co-immunoprecipitation (Co-IP) assay in *N. benthamiana* leaves expressing GFP-tagged SCP8 and myc-tagged NbSBT1 (Fig. 3D). To assess whether SBT activity is involved in SCP8 action, we co-expressed two SBT inhibitors, SPI-1 from Arabidopsis^40^ and EPI1 from oomycete pathogen *Phytophthora infestans*^41^ with SCP8 (or free GFP as the control) in *N. benthamiana*, followed by measurement of the ROS burst triggered by different doses of flg22. As shown in Fig. 3E, co-expression of SCP8 with SBT inhibitor SPI-1 or EPI1 significantly reduced SCP8-mediated defense suppression. From these findings we can conclude that SBT activity is required for SCP8-mediated plant immune suppression. It is conceivable that SCP8 may have activated or enhanced subtilase activity, generating more phytocytokines including PSK, resulting in PTI suppression.

Consistent with a role of PSK in SCP8-mediated immune suppression, flg22-but not chitin-induced ethylene production in Col-0 was strongly suppressed by concomitant infiltration of PSK. Likewise, ethylene production was restored in the PSK receptor mutant *pskr1-3* (Fig. 4A). To confirm a requirement of PSK signalling for SCP8 action, we attempted to express *SCP8* in Arabidopsis PSK-receptor mutant genotypes *pskr1-3, pskr1-3/pskr2*, and *pskr1/psy1r*. Unfortunately, for unknown reasons, we could not obtain a single stable transgenic line, suggesting that the constant presence of the effector impairs essential plant functions. Alternatively, apoplastic fluids of *N. benthamiana* plants expressing SCP8-GFP and free GFP were harvested (Fig. S3A) and infiltrated together with flg22 or pg9 into Col-0 and *pskr1-3* leaves. As expected, Col-0 leaves co-infiltrated with apoplastic fluid containing SCP8 and flg22 or pg9, respectively, showed more than 50% lower ethylene production compared to that observed upon co-infiltration of apoplastic fluid containing GFP and any of the two patterns. Notably, ethylene production was partially restored in leaves of the *pskr1-3* mutant when treated simultaneously with SCP-containing apoplastic fluid and elicitor (Fig. 4B). We repeated this experiment with recombinant His-tagged SCP8 protein produced in *Pichia pastoris* (Fig. S3B), which was infiltrated into Arabidopsis leaves prior to pattern treatment. In Col-0, leaves pretreated with SCP8 showed an approximately 30% reduction in ethylene production in response to flg22 or pg9 compared to leaves pretreated with water. In contrast, in leaves of *pskr1-3* plants, the loss of pattern-induced ethylene production caused by SCP8 was fully restored (Fig. 4C). Taken together, our findings suggest that SCP8 utilizes PSK signaling to dampen host immune activation.

**Fig. 4.**
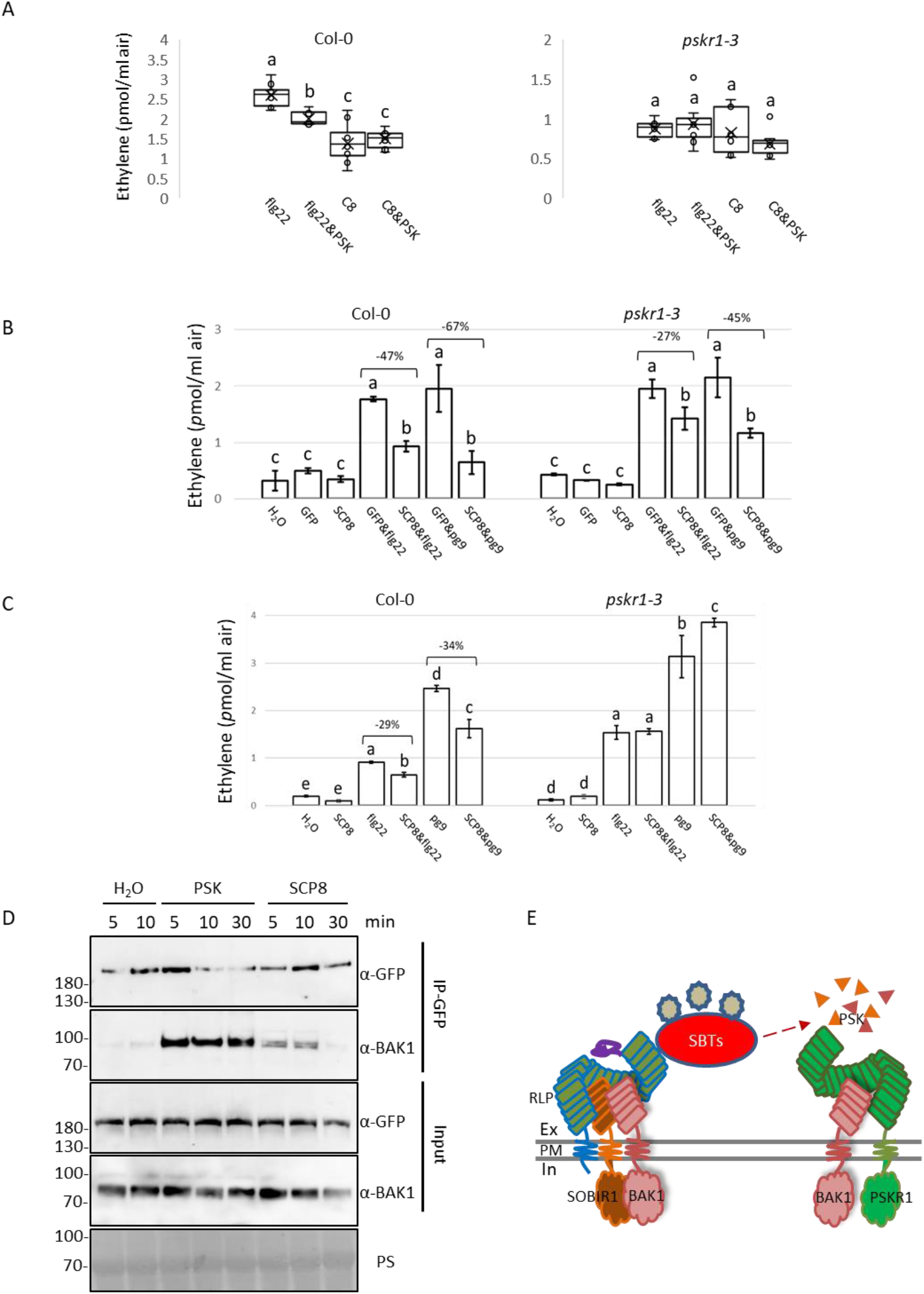
SCP8 suppresses PTI by activating PSK signaling. A, Ethylene production of Col-0 and *pskr1-3* mutant 2 hours after infiltration of flg22 (1 µM), flg22&PSK (both are 1 µM), C8 (1 µM) and C8&PSK (both are 1 µM), respectively. B, Ethylene production of Col-0 and *pskr1-3* 2 hours after infiltration of H_2_O, *N. benthamiana* apoplastic fluids containing GFP or SCP8 (hereafter GFP or SCP8), GFP&flg22 (1 µM), GFP&pg9 (1 µM), SCP8&flg22 (1 µM), and SCP8&pg9 (1 µM). C, Ethylene production of Col-0 and *pskr1-3* infiltrated with H_2_O or SCP8 protein prior to flg22 (1 µM) and pg9 (1 µM) treatments. D, Western blot of Co-IP samples from transgenic Arabidopsis expressing PSKR1-GFP infiltrated with H_2_O, 1 µM PSK and 1 µM SCP8, respectively. Input and GFP-trap beads precipitated samples were detected by both anti-GFP and anti-BAK1 antibodies. The different letters depict statistically significant differences (P<0.05, student’s t-test). E, The model of SCP8 function.

PSK activates the formation of PSKR1-BAK1 complexes^14^, suppressing PTI^9-11^. To test whether SCP8 activates the formation of PSKR1-BAK1 complexes, Arabidopsis plants stably expressing *p35S::AtPSKR1-GFP* were infiltrated with recombinant SCP8 and subsequently subjected to immunoprecipitation by GFP-trap agarose beads. Co-Immunoprecipitation of *At*BAK1 along with *At*PSKR 1-GFP was observed at 5 and 10 minutes after SCP8 infiltration, but not after treatment with water as negative control (Fig. 4D). Likewise, treatment with PSK, which was used as positive control, resulted in precipitation of *At*BAK1 levels that were even higher than those observed in plants treated with SCP8. While we can only speculate about the reason for this difference we assume that PSK concentrations produced upon SCP8 treatment may differ significantly from those used for PSK treatment in our experiment (Fig. 4D). A similar experiment was also performed in *N. benthamiana* transiently co-expressing *p35S::AtPSKR1-GFP* and *p35S::AtBAK1-myc* constructs. Consistent with our findings in Arabidopsis, SCP8 also activated the formation of *At*PSKR1-*At*BAK1 complexes in a heterologous plant (Fig. S2).

PSK-PSKR1 signaling has previously been shown to negatively regulate plant innate immunity, likely due to the recruitment of the PRR co-receptor BAK1, which then restricts the availability of BAK1 for the formation of PRR-BAK1 complexes that are required for PTI activation^14^. We here show that a multi-host fungal plant pathogen utilizes a secreted protein effector to manipulate PTI activation by enhancing signaling of an immunosuppressive phytocytokine, PSK (Fig 4E). SCP8 is one of the few known microbial effectors that target an apoplastic factor to suppress PTI ^1,42^. Effector-mediated manipulation of a negative immunoregulatory phytocytokine contributes to a bewildering array of immunosuppressive infection strategies of plant microbial pathogens.

## Methods

### Fungal strains and transformation

Cotton isolated *Verticillium dahliae* (*Vd*) strain 592 (V592)^43^ was used as wildtype strain and recipient strain for DNA transformation. *Agrobacterium tumefaciens* strain EHA105 was used for *Vd* transformation. Potato dextrose agar (PDA) medium with or without antibiotics was used to culture V592 and transformants as described previously^33^.

### Genes and constructs

The VdSCP8 gene knockout construct was made following the strategy of Wang *et al*. ^33^with primers listed in Table S3. SCP8 gene construct for genetic complementation was made as described^27^ with primers listed in Table S3. For the VdSCP8-GFP fusion construct, the Olic promoter and open reading frames of VdSCP8 and GFP were amplified by PCR (primers in Table S3), and then joined by overlap extension PCR with primers PoliC_pSul_F and GFP_pSulC_R (Table S3). The product was cloned into pSul-RG#PB vector^33^ digested with HindIII and EcoRI using ClonExpress II One-Step Cloning Kit (Vazyme, China).All the constructs for *in planta* expression were made in pLOCGex vector with primers listed in Table S3 following the method of Zhang *et al*. ^27^

### Arabidopsis transformation and selection

*Arabidopsis thaliana* Col-0 was transformed by *A. tumefaciens* GV3101 strain expressing pLOCGex::SCP8, and T0 seeds were selected on ½ MS plates containing 50 µg/ml kanamycin.

### Transient expression in *Nicotiana benthamiana*

*A. tumefaciens* strains were grown in YEB medium (0.5% (w/v) beef extract, 0.5% (w/v) bacteriological peptone, 0.5% (w/v) sucrose, 0.1% (w/v) yeast extract, 2 mM MgSO4) with appropriate antibiotics and 20 µM acetosyringone at 28 °C for overnight. Cultures were harvested and resuspended in 2.0% (w/v) sucrose, 0.5% (w/v) Murashige and Skoog salts without vitamins, 0.2% (w/v) MES, and 0.2 mM acetosyringone, pH 5.6, to an optical density at 600 nm (OD_600_) = 1.0. For co-expression, two cultures carrying appropriate constructs were mixed in a 1:1 ratio to OD600 = 0.5 for each. The resuspended cultures were incubated at room temperature for 1-3 h, and infiltrated into 5-6-week-old *N. benthamiana* leaves with 1 ml syringe. Samples were collected 1 or 2 days after agro-infiltration for gene expression analysis, microscopic analysis or immunoblotting analysis. Agro-infiltration was performed at least 2 times for each analysis.

### RNA extraction and reverse transcription quantitative PCR (RT-qPCR)

Total RNA was isolated using a Nucleospin® RNA Plant Kit (Macherey-Nagel, Germany) according to the manufacturer’s instructions. First-strand cDNA was synthesized from 1 µg of total RNA with SuperScript® III Reverse Transcriptase (Invitrogen), according to the manufacturer’s instructions. RT-qPCR was performed using a C1000 Thermal Cycler (Bio-Rad) in combination with the EvaGreen 2x qPCR MasterMix - iCycler (Applied Biological Materials). The primers used to detect the transcripts are listed in Table S1. The RT-qPCR conditions were as follows: an initial 95 °C denaturation step for 3 min, followed by denaturation for 15 s at 95 °C, annealing for 20 s at 60 °C, and extension for 20 s at 72 °C for 40 cycles. The data were collected using the Bio-Rad CFX Manager software (Bio-Rad, US) and processed in Microsoft Excel. The transcript levels of target genes were normalized to the transcript levels of the constitutively expressed *VdGAPDH* gene^44^ or *NbEF1α* gene^45^ (Table S3) according to the 2^-ΔΔCt^ method^46^.

### Elicitors and immune response measurements

Chitin (IsoSep, No.57/12-0010, MW 1643.57) was dissolved in water to make 100 µM stock. flg22, nlp20, and pg9 were synthesized by Genscript and dissolved in water to make 100 µM stocks. Elicitor induced immune responses including ROS burst, MAPK activation, and ethylene production were measured following the methods described previously^17^. To measure ethylene production after protein or peptide infiltration, the elicitor was added to MES buffer (pH 5.6) with infiltrated leaf pieces, followed by ethylene measurement after 2 hours.

### Protein purification from *Pichia pastoris*

The SCP8 gene was amplified by PCR with primers pPSCP8_F and pPSCP8_R (Table S3), and ligated into pPICZα vector digested by *EcoR*I and *Not*I restriction enzymes, yielding pPICZα::SCP8 construct. The construct was transformed into *P. pastoris* GS115 strain, and transformants were selected on YPD medium containing 50 µg/ml zeiocin. Single colony of SCP8 transformant was inoculated into YPD medium, and protein expression was induced by methanol in BMMY medium following the description in Pichia protein expression handbook (Invitrogen). SCP8 protein was purified by affinity chromatography with a His-trap column (Cytiva) followed by size exclusion chromatography with Superdex™ 200 column (Cytiva).

## Data availability

RNAseq data (NCBI/GEO/upload/chenleihua2017/_qFOZaGag)

## Acknowledgements

This work was supported by Deutsche Forschungsgemeinschaft (DFG) grant Nu70/17-1 to T.N., the National Natural Science Foundation of China (No. 32020103003) to H.-S.G, and National Natural Science Foundation of China (31500119) and Visiting Scientist Fellowship (201604910325) of China Scholarship Council to C.H.

## Author contributions

C.H., H.-S.G., and T.N. conceived and designed major experiments; A.S., A.S. and C.H. designed the experiments with protease inhibitors. L.Z. and C.H. conducted experiments; C.H. and T.N. wrote the manuscript. All authors analyzed data, discussed the results and commented on the manuscript.

## Additional information

Supplementary files include Fig. S1-S3 and Table S1-S3.

**Fig. S1.**
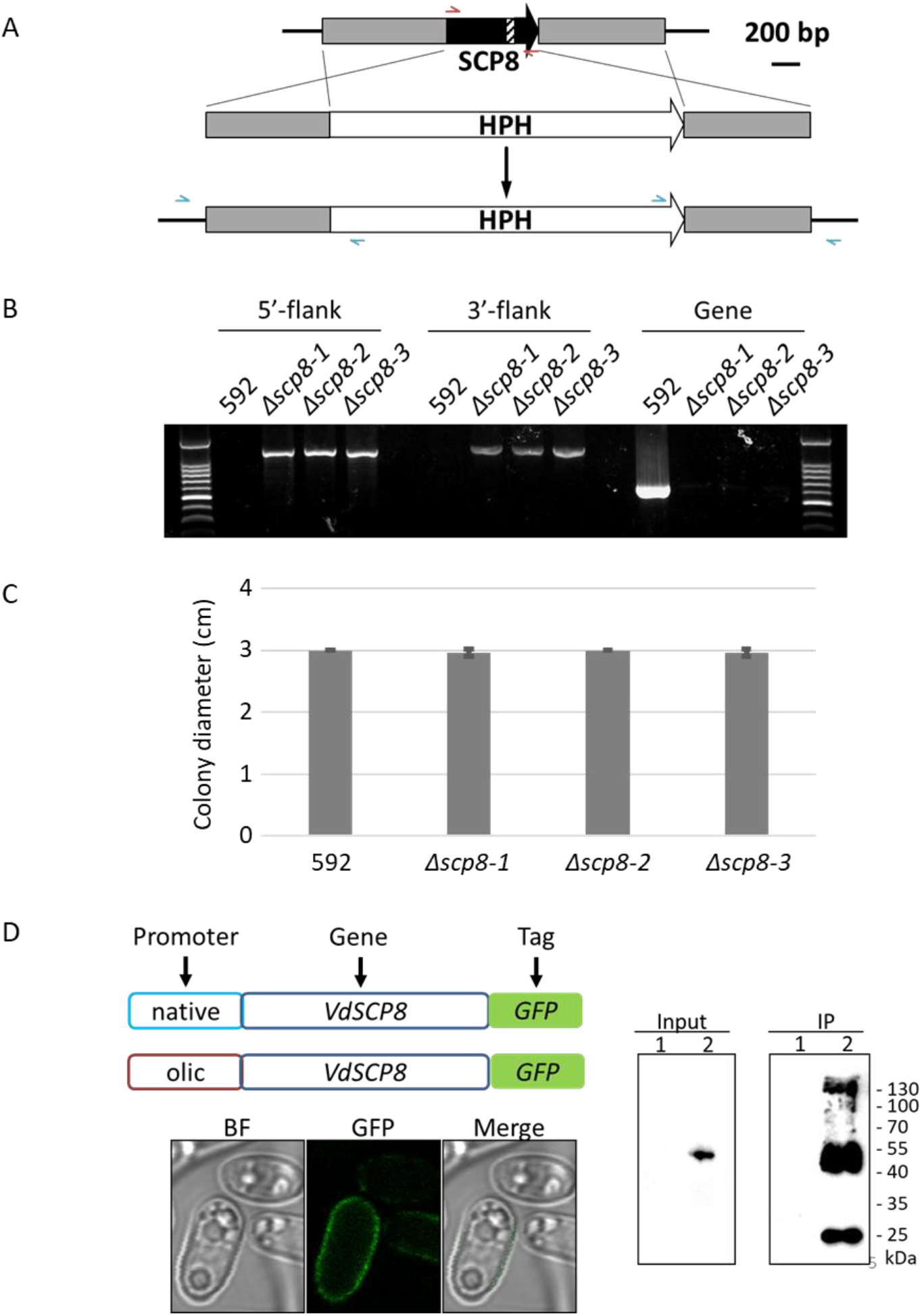
VdSCP8 gene knockout and complemented transformants. A, SCP8 gene knockout strategy. B, Verification of SCP8 gene knockout mutants by genomic PCR. C, Colony diameter of *Vd* wild-type strain (592) and SCP8 knockout mutants *Δscp8-1, Δscp8-2, Δscp8-3*. D, Verification of SCP8 complemented transformants by observation of GFP fluorescence and GFP antibody (1, *Vd* wild-type, 2, C-olic strain).

**Fig. S2.**
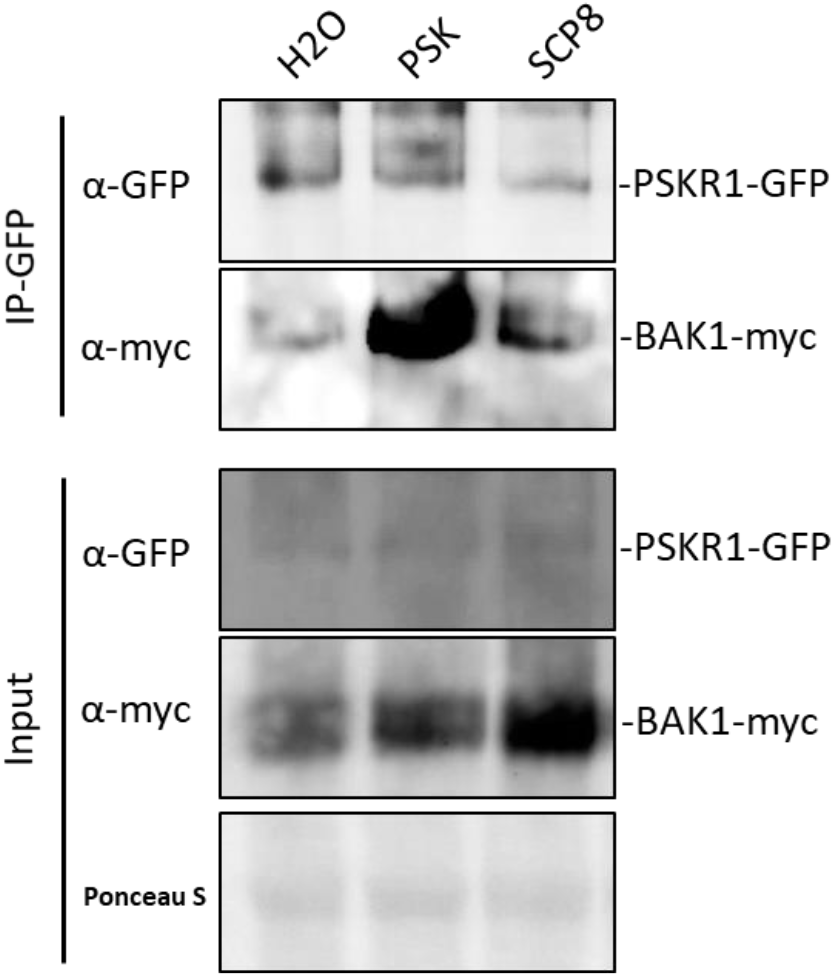
Western blot of Co-IP samples from *N. benthamiana* co-expressing PSKR1-GFP and BAK1-myc infiltrated with H_2_O, 1 µM PSK and 2 µM SCP8, respectively. Input and GFP-trap beads precipitated samples were detected by both anti-GFP and anti-myc antibodies.

**Fig. S3.**
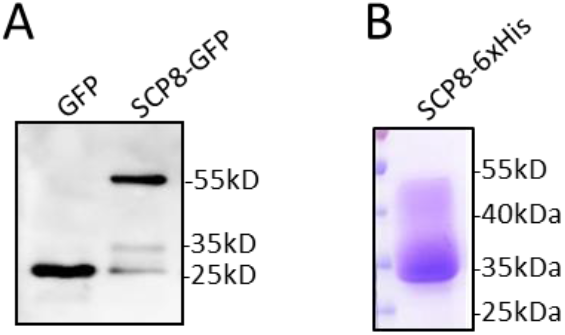
SCP8 expressed in *N. benthamiana* and *Pichia pastoris*. A, Apoplastic fluids isolated from *N. benthamiana* transiently expressing GFP or SCP8-GFP, and detected by GFP antibody. B, C-terminal 6xHis-tagged SCP8 fusion protein isolated from *Pichia pastoris* on SDS gel stained by coomassie brilliant blue.

